# Identifying the best PCR enzyme for library amplification in NGS

**DOI:** 10.1101/2022.10.31.514486

**Authors:** Michael A Quail, Craig Corton, James Uphill, Jacqueline Keane, Yong Gu

## Abstract

**Background:** PCR amplification is a necessary step in many next generation sequencing (NGS) library preparation methods[1] [2]. Whilst many PCR enzymes are developed to amplify single targets efficiently, accurately and with specificity, few are developed to meet the challenges imposed by NGS PCR, namely unbiased amplification of a wide range of different sizes and GC content. As a result PCR amplification during NGS library prep often results in bias toward GC neutral and smaller fragments. As NGS has matured, optimised NGS library prep kits and polymerase formulations have emerged and in this study we have tested a wide selection of available enzymes for both short read Illumina library preparation and long fragment amplification ahead of long-read sequencing.

**Results:** We tested over 20 different Hi-fidelity PCR enzymes/NGS amplification mixes on a range of Illumina library templates of varying GC content and composition, and find that both yield and genome coverage uniformity characteristics of the commercially available enzymes varied dramatically. Three enzymes Quantabio RepliQa Hifi Toughmix, Watchmaker Library Amplification Hot Start Master Mix (2X) “Equinox” and Takara Ex Premier were found to give a consistent performance, over all genomes, that mirrored closely that observed for PCR free datasets. We also test a range of enzymes for long read sequencing by amplifying size fractionated S. cerevisiae DNA of average size 21.6 and 13.4kb respectively.

**Conclusion:** The enzymes of choice for short read (Illumina) library fragment amplification are Quantabio RepliQa Hifi Toughmix, Watchmaker Library Amplification Hot Start Master Mix (2X) “Equinox” and Takara Ex Premier, with RepliQa also being the best performing enzyme from the enzymes tested for long fragment amplification prior to long read sequencing.

## Background

In barely over a decade Next Generation Sequencing (NGS) has transformed the biological sciences and is now used for a diverse range of applications on a diverse range of species and sample types. From the early days of NGS the concept of bias was recognised. No existing sequencing technology can “read” a genome from start to end so DNA/RNA are fragmented to a size that can be read by the sequencing platform, a library of those fragments is prepared and original sample sequence is reconstituted following sequencing of the resulting mixture of library fragments. Ideally all of the constituent parts of the genome would be sequenced with equal representation, though in practice this is never the case. During the library prep and sequencing process there are several points that can introduce a bias into the representation, though perhaps the single most-bias introducing step is PCR. Sequencing libraries are comprised of a mixture of fragments of different size, GC and repeat content that together represent the original sample. Smaller, more GC neutral, fragments that don’t contain any secondary structure will amplify more efficiently than larger, high GC, high AT fragments and those that are capable of forming secondary structures. Over multiple cycles any bias will be amplified so the number of PCR cycles should be kept to a minimum, indeed for the most even coverage sequencing dataset PCRfree libraries are recommended [3]. Whilst advantageous, PCRfree methods are not always practicable due to the high DNA mass input requirements, necessitating the use of PCR amplification during library prep.

Since its invention in 1986 PCR has proven to be a convenient and effective way to selectively amplify genomic loci of interest and is the most common approach used for targeted selection ahead of both short and long-read sequencing. Whilst the maximal fragment size sequenceable in a contiguous manner using short read sequencing is around 600bp, Pacific Bioscience and Oxford Nanopore Technologies sequencing platforms can generate reads in excess of 100kb. This capability has led to long read sequencing approaches being preferred for *de-novo* sequencing. However, the long-read technologies require significant amounts of input DNA meaning amplification during long read library preparation may be necessary. Here we compare enzymes for their ability to amplify long DNA fragments and compare yield and sequencing performance.

In 2011 we published a study that identified Kapa HiFi as the best enzyme for Illumina library amplification steps as it gave the most even coverage across a set of microbial genomes of diverse GC content [4]. Whilst most enzymes gave relatively even coverage over the more GC neutral genomes used (*Salmonella Pullorum* and *Staphylococcus aureus*) large differences in coverage representation were observed with libraries generated using different enzymes for the GC rich genome of *Bordetella pertussis* and the AT rich genome of *Plasmodium falciparum*.

Ten years on, we perform a similar study using many currently available and newly developed High Fidelity polymerase mixes/NGS amplification formulations. There are a great many commercially available PCR enzymes. Enzymes were chosen for this evaluation through discussion with individual enzyme manufacturers and suppliers to identify their enzymes that they recommended for NGS, these were mostly hotstart formulations of type II enzymes that have been developed to amplify complex templates and introduce fewer errors than standard Taq polymerases.

## Results

### Initial evaluation of enzymes for Illumina library amplification

Each enzyme was assessed for its ability to amplify genomic Illumina adapter ligated library fragments of an expected average insert size of approximately 500bp, from a set of four microbial genomes with differing GC content:- *Bordetella pertussis*, 67.7 % GC; *Escherichia coli,* 50.8 % GC; *Clostridioides difficile*, 29.1 % GC; and *Plasmodium falciparum,* 19.3 % GC. For each enzyme tested, 1 ng of pre-PCR library fragments from each genome was amplified using manufacturers recommended denaturation and extension times, with annealing at 60°C for 15 seconds and 14 cycles. Unique dual indexed P7 and P5 amplification primers were used to avoid index hopping [5, 6].

For UDI oligonucleotides used, see **Supplementary Table 1.**

Details of enzymes used and cycling conditions are listed in **Supplementary Table 2**.

After 0.7x Ampure bead cleanup the yield of each library was assessed using fluorimetric measurement (**Supplementary Figure 1**). There were surprising differences in the yields obtained from the enzymes tested with some giving relatively little library. These were repeated with fresh enzyme on a different PCR block, always with the same outcome. Quantabio RepliQa and sparQ, Kapa HiFi, Invitrogen Platinum Superfi II, Thermo Collibri and Phusion U multiplex PCR mastermix, Tools Ultra, Biotool Univerase and Agilent Herculase gave good yields.

Barcoded libraries were pooled in a pseudoequimolar manner according to genome size and run on an Illumina Novaseq 6000 SP or S4 flowcell lane to give > 30x coverage of each genome. To fairly compare results, datasets were randomly trimmed to contain reads representing 30 x coverage. We tabulated the depth of coverage seen at each position of the genome and calculated the fraction of each genome (referred to as Low Coverage Index) that was covered to a depth of less than 15 x i.e. half the mean coverage. Most datasets had <5% low coverage with the GC neutral *E. coli* genome but higher degrees of low coverage were observed for less base balanced genomes with a lot of enzyme, with the extremely AT rich genome of *P falciparum* posing the biggest challenge (**Figure 1**). Quantabio RepliQa, Kapa HiFi, and Collibri had <5% low coverage index with all four genomes.

**Figure 1.**
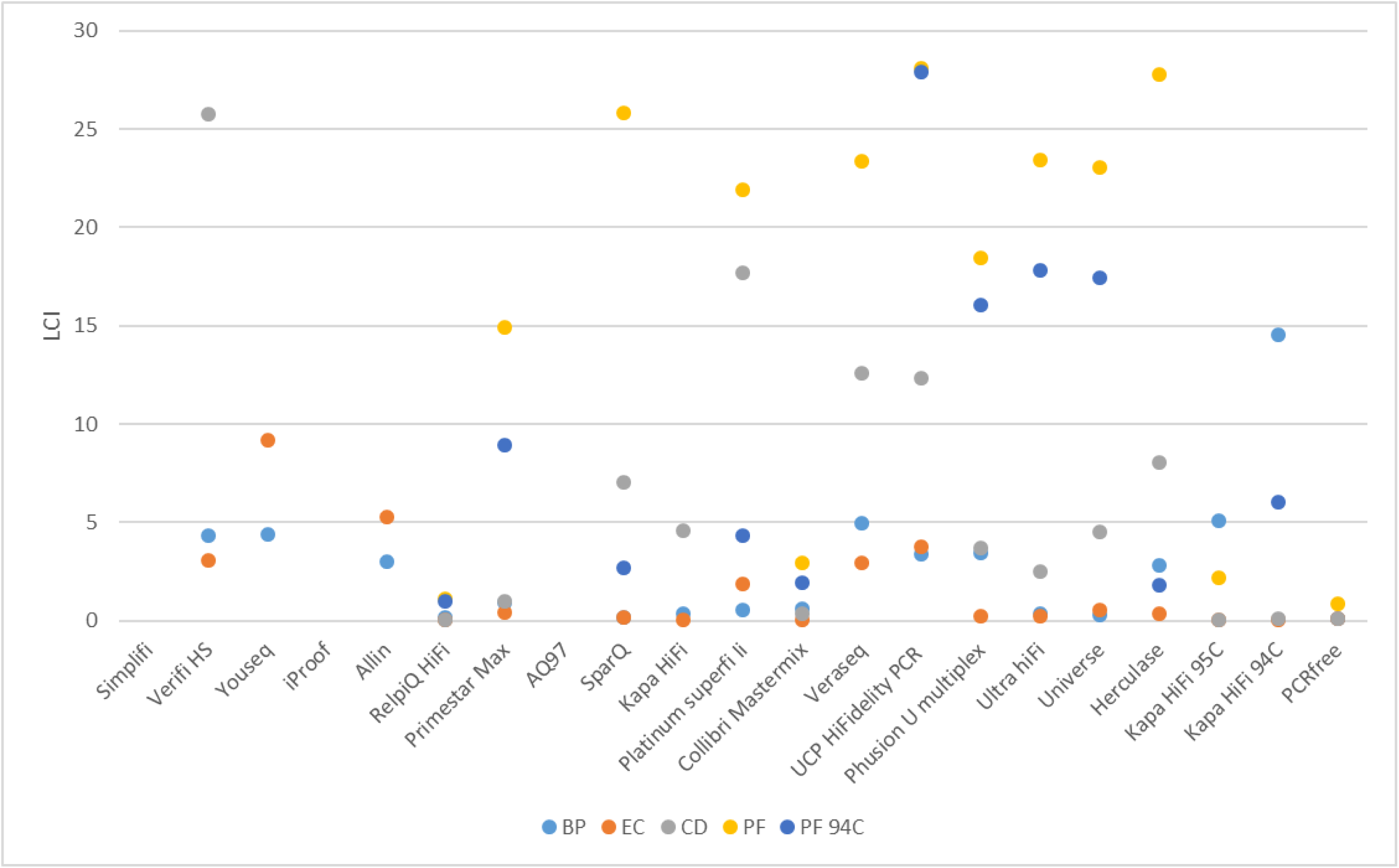
Low coverage index obtained after 14 cycles of amplification with 1ng of each test microbial genome; BP: *Bordetella pertussis*, EC: *Escherichia coli*, CD: *Clostridioides difficile*, and PF: *Plasmodium falciparum*. For each PF was also amplified using a 94C denaturation temperature, “PF 94C”.

### Further evaluation of enzymes for Illumina library amplification

To test reproducibility further replicate libraries were made from the better performing enzymes, with the addition of Watchmaker Genomics Equinox library amplification mastermix, and Takara Ex Premier (these new formulations had been previously unavailable for testing), under a variety of different cycling conditions; **Supplementary Table 2**. With this selected group, yields were quite high with all templates (**Supplementary Figure 2 and 3**).

Again the low coverage index was calculated for each dataset and enzymes/conditions ranked from low to high LCI for each genome (**Figure 2**).

**Figure 2.**
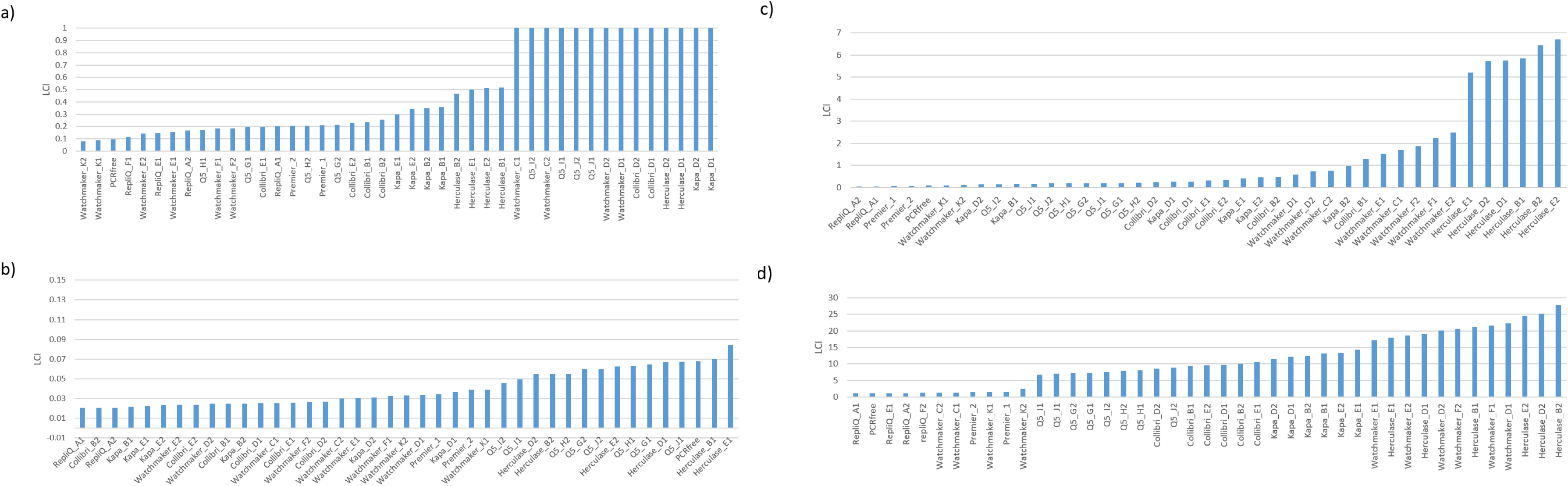
Ranked Low Coverage Index values obtained after 14 cycles of amplification with 1ng of each test microbial genome; a) *Bordetella pertussis*, b) *Escherichia coli*, c) *Clostridioides difficile*, and d) *Plasmodium falciparum*.

Whilst some enzymes perform better in certain genomic contexts RepliQa, Watchmaker Equinox, and Takara Ex Premier, give good coverage uniformity with all genomes. To assess the average low coverage index across all genomes we calculated the sum of the low coverage values compared to coverage from PCRfree libraries (**Figure 3**) illustrating that these three enzymes have minimal bias each giving coverage uniformity similar to that seen with PCR free libraries.

**Figure 3.**
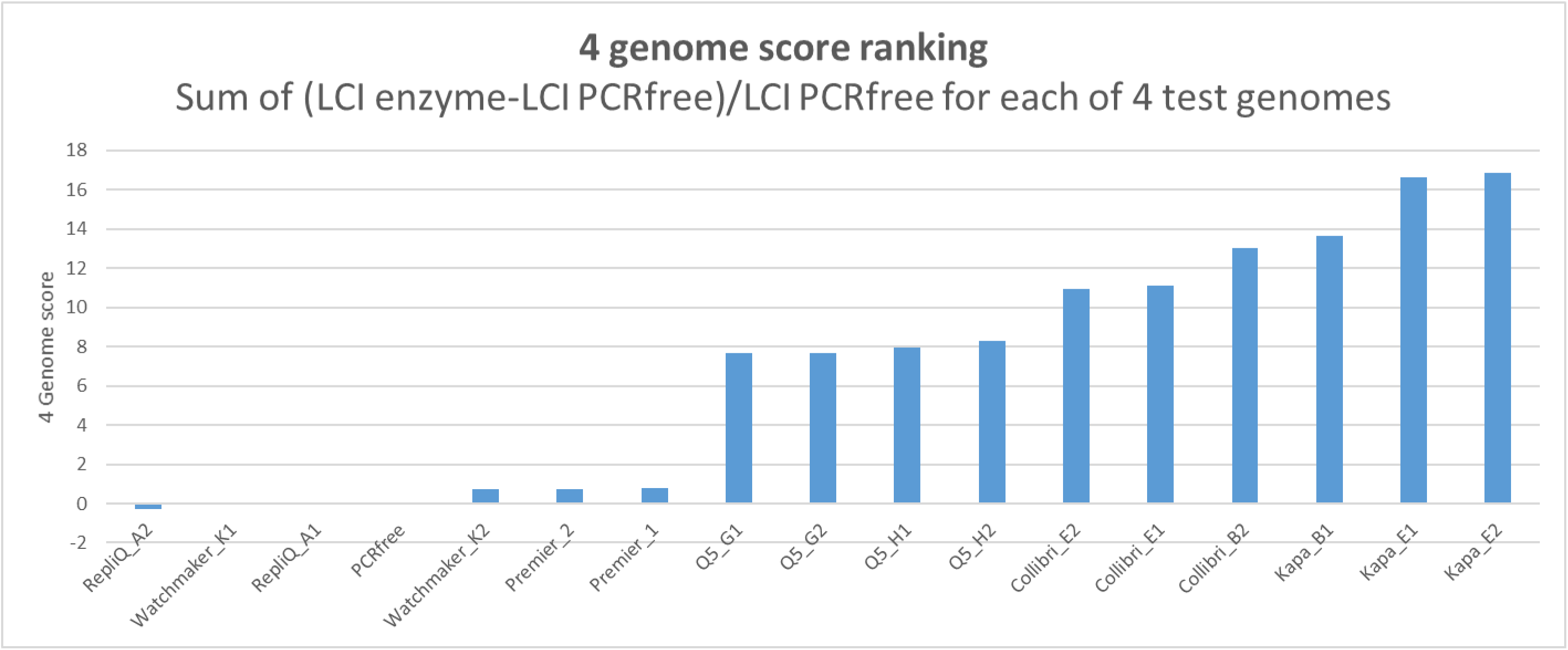
Multi-genome average low coverage index ranking taking the sum of the LCI values for all 4 genomes compared to that obtained from PCR free data.

The end result of the more even coverage obtained with RepliQa and Equinox relative to other enzymes could be clearly seen in the more challenging GC or AT rich regions where RepliQa, Equinox and PCR free had good coverage in GC rich (locally 100% GC) regions of *B. pertussis* and also in AT rich (locally <4% GC) regions of *P falciparum* (**Figure 4**).

**Figure 4.**
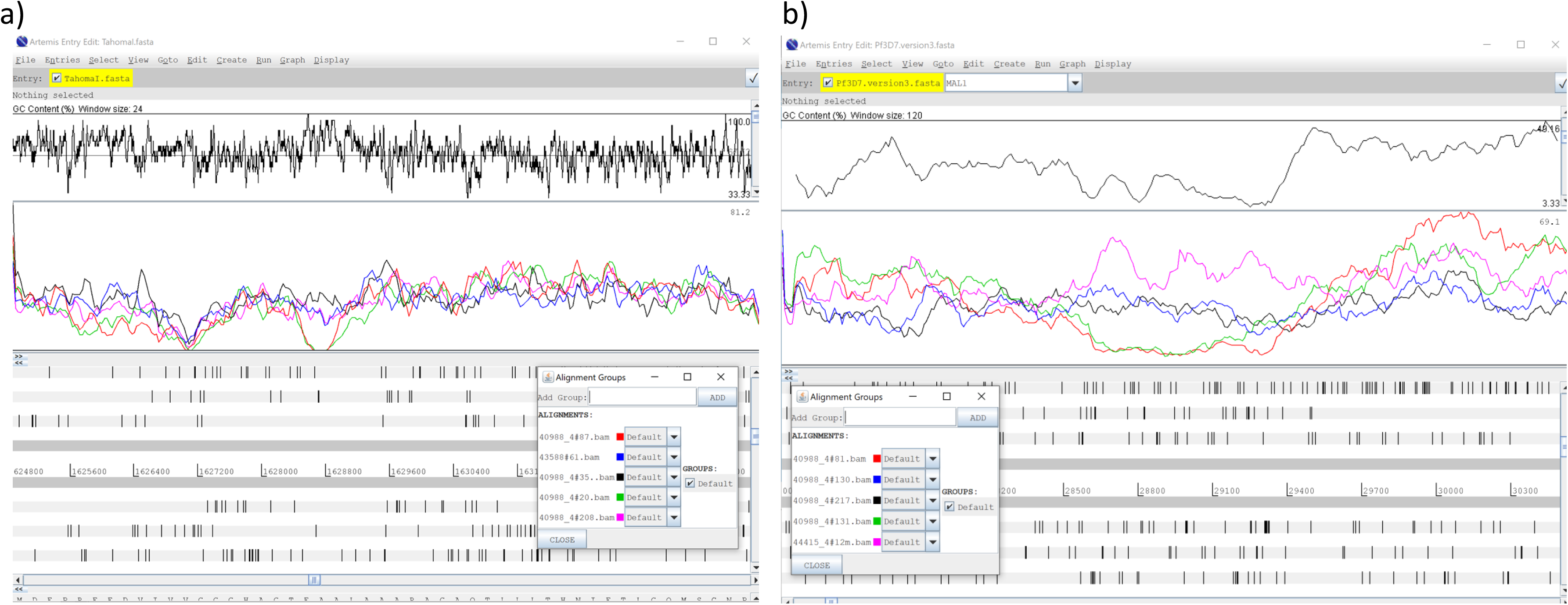
Screenshot from Artemis genome browser [7] over regions of a) *B. pertussis* and b) *P. falciparum* genomes. In each the top panel is a GC content plot with the maxima and minima GC content of each region displayed at the far right. The bottom panel is an open read frame plot and the middle panel plots read coverage over each base. The inset panel on each denotes the enzyme used for each coloured line. In a) RepliQa (pink), Watchmaker (blue) and PCRfree data (black) coverage is unaffected whereas with kapa hifi (red) and Collibri (green) coverage drops to near zero in the GC rich region in the region in the centre of the plot. Likewise in b) RepliQa (blue), Watchmaker (black) and PCRfree data (pink) coverage is unaffected whereas with kapa hifi (red) and Collibri (green) coverage drops to near zero in the AT rich region in the region in the centre of the plot.

Sequencing data was also obtained for human genome template amplification. Again this showed RepliQa, Watchmaker Equinox and Takara Ex Premier to have the most even genome coverage reflected in the lowest LCI values (**Supplementary Figure 4**).

It has been observed that some enzymes are inhibited with magnetic beads that are commonly used in NGS workflows e.g. SPRI magnetic bead cleanup and size selection, and streptavidin bead capture of biotin labelled DNA fragments. With SPRI cleanup carryover of beads after elution is commonplace and some protocols employ “with bead” approaches where increased yield is obtained when the beads are not removed after the final elution step [7]. Streptavidin conjugated magnetic beads are also used in NGS protocols for selection of biotin labelled library fragments, most commonly in hybrid capture target enrichment procedures [8], and due to the extremely strong affinity of streptavidin for biotin such methods require PCR amplification of bead bound library fragments. To test if amplification by the enzymes used in the second phase of this study are inhibited by such beads amplification was carried out without beads, with a volume of washed beads in water equivalent to equal sample volume or with the template bound to 50ul of washed streptavidin beads (Dynabeads MyOne Streptavidin T1, Thermo, cat no. 65602). All of these enzymes tested were found to be unaffected by the presence of Ampure or streptavidin magnetic beads, apart from Q5 (**Supplementary Figure 5**).

Accuracy and utility of the human genome sequences obtained with each enzyme were assessed by comparing each dataset with the NA12878 reference genome and variant list (see methods). Both numbers of indels and SNPs detected were slightly greater with 500bp mean inserts compared to 200bp. The three enzymes with the highest sensitivity for SNP and indel detection in the microbial reference genomes were QuantaBio RepliQa, Watchmaker Equinox and Takara Ex Premier (**Table 1**). These enzymes were found to call more SNPs and indels with greater precision, compared to kapa HiFi at rates that are comparable to those seen in the Precision FDA Truth challenge when using 50x PCRfree datasets [9].

**Table 1.** Indel True Positive, False Negative and Sensitivity values for NA12878 libraries prepared with either approx 200 or 500 bp inserts with different PCR enzymes.

| Sample ID | No. SNPs (TP) | No. SNPs (FN) | SNP Sensitivity (%) | No. Indels (TP) | No. Indels (FN) | Indel Sensitivity (%) |
| --- | --- | --- | --- | --- | --- | --- |
| repliQ_200bp | 3264914 | 93588 | 97.21 | 468524 | 67965 | 87.33 |
| kapa HiFi_200bp | 3271956 | 86546 | 97.42 | 457018 | 79471 | 83.86 |
| kapa HiFi_500bp | 3322273 | 36229 | 98.92 | 470902 | 65587 | 85.19 |
| herculase_500bp | 3293130 | 65372 | 98.05 | 464692 | 71797 | 86.62 |
| repliQ_500bp | 3333633 | 24869 | 99.26 | 502494 | 33995 | 93.66 |
| Watchmaker_500bp | 3333145 | 25357 | 99.25 | 490866 | 45623 | 91.5 |
| Premier_500bp | 3330695 | 27807 | 99.17 | 503501 | 32988 | 93.85 |

There may be times in a high throughput lab or within a clinical sequencing lab when fast turnaround is required when rapid PCR may be desired. Quantabio promote RepliQa for its short extension times. When we tested the enzymes in phase 2 with increasing extension times (5, 15, 30 or 60 seconds) it was observed that Collibri, Q5, RepliQa and Watchmaker Equinox enzymes gave near maximal yield after just 5 seconds of extension whereas Herculase and Kapa HiFi yields increased with extension time (**Supplementary Figure 6**).

The genome of the malaria parasite *Plasmodium falciparum* has an extremely low GC content of 19.3% [10] and has been shown to be one of the most challenging genomes to amplify and sequence [3, 4] [11]. There have been several papers published for this genome detailing methods to minimise the biases introduced by PCR and sequencing including PCR free library approaches [3], [12] and optimised PCR protocols [4, 11] [13, 14]. In this study when using these approaches we find that the fraction of the genome at less than 50% of mean coverage could be decreased even further (**Figure 5**) though the most successful reduction was achieved by using a different approach for different enzymes. The lowest LCI was achieved using RepliQa with denaturation at 94C and extension at 60C as described by Lopez-Barragan, though near similar LCI values were also obtained under these conditions using Collibri and Watchmaker Equinox enzymes. Following on from this we tested RepliQa under a range of denaturation and extension temperature combinations and found that these conditions could not be improved upon (results not shown).

**Figure 5.**
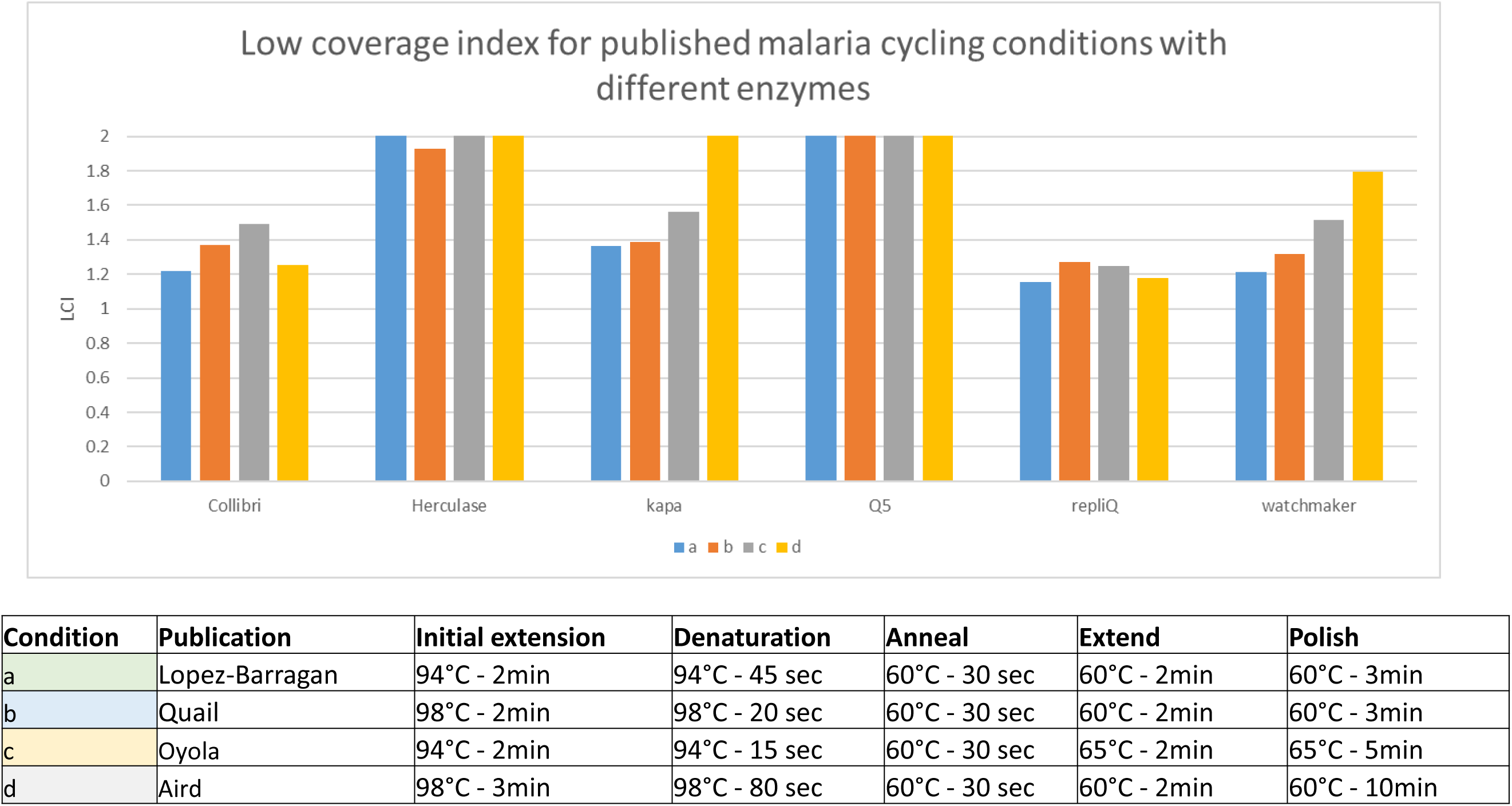
Low coverage index for amplification of 1ng P falciparum Illumina library template with a range of enzymes using PCR conditions described by Lopez-Barragan [14] (blue), Quail (orange) [4], Oyola [12] (grey) and Aird [15] (yellow).

### Long range amplification for Long Read sequencing

Long range PCR is a common approach for generation of material for long read sequencing. Many users have found this to be even more challenging with low yield and a bias towards smaller fragments during amplification. To test the suitability of PCR enzymes for this application we prepared size fractionated adapter ligated yeast genome fragments adding Illumina adapters to enable amplification using the same primers as used in the rest of this study.

Sheared *S. cerevisiae* DNA was size fractionated using Sage Sciences ELF or Bluepippin instruments yielding modal fragment sizes of 21.6kb and 13.3kb respectively (**Supplementary Figure 7**). After adapter ligation 1ng of each of these were used as a template for long range PCR with a range of enzymes using manufacturers recommended cycling conditions (**Supplementary Table 2**).

Initially, 12 cycles of PCR was used, but with most enzymes that generated little or no product (data not shown) so PCR was repeated for 15 cycles after which time amplicons of the expected size were observed with most enzymes (**Supplementary Figure 8**), though yields varied widely (**Supplementary Table 3**). The long range PCR products were then prepared for Pacific Biosciences HiFi sequencing using manufacturers’ recommended amplicon library prep protocol and barcoded adapters. Sequencing yields and coverage obtained are summarised in (**Supplementary Figure 9 and Table 2)**. Due to extremely low yields after PCR, products from some enzymes gave insufficient yield to obtain significant coverage.

**Table 2.** Assembly statistics from PacBio HiFI 30 x genome coverage from the various PCR enzyme datasets.

| Downsampled to 30x |  |  |  |  |  |  |  |  |  |  |  |  |  |
| --- | --- | --- | --- | --- | --- | --- | --- | --- | --- | --- | --- | --- | --- |
| Enzyme | Template | PCR cycles | Reads | Yield | Fold Coverage | Sample Name | Polished Contigs | Maximum Contig Length | Mean Contig Length | Median Contig Length | N50 Contig Length | Sum of Contig Lengths | Number of Circular Contigs |
| repliQa | ELF | 15 | 95,403 | 874,550,481 | 72.88 | repliQAx15 | 20 | 1,529,563 | 609,561 | 623,779 | 834,867 | 12,191,225 | 2 |
| repliQa | Bluepippin | 15 | 67,874 | 851,073,366 | 70.92 | repliQaBx15 | 22 | 1,092,081 | 551,212 | 597,022 | 746,483 | 12,126,668 | 2 |
| repliQa | Bluepippin | 12 | 39,211 | 508,550,579 | 42.38 | repliQaBx12 | 22 | 1,533,220 | 553,116 | 586,171 | 784,885 | 12,168,568 | 2 |
| Terra | Bluepippin | 15 | 69,375 | 811,670,287 | 67.64 | terraBx15 | 23 | 1,097,683 | 528,785 | 566,757 | 785,063 | 12,162,064 | 3 |
| Terra | ELF | 15 | 132,515 | 851,869,210 | 70.99 | terraAx15 | 33 | 1,092,332 | 368,810 | 339,580 | 509,984 | 12,170,747 | 2 |
| Q5 | Bluepippin | 15 | 76,603 | 523,297,797 | 43.61 | Q5Bx15 | 35 | 880,290 | 345,193 | 274,664 | 474,198 | 12,081,766 | 2 |
| LongAmp | Bluepippin | 15 | 69,634 | 626,495,361 | 52.21 | longampBx15 | 39 | 934,826 | 310,307 | 283,489 | 499,795 | 12,102,002 | 1 |
| Universe | Bluepippin | 15 | 51,436 | 470,686,450 | 39.22 | universeBx15 | 104 | 604,982 | 111,981 | 89,487 | 161,100 | 11,646,056 | 2 |
| Watchmaker Equinox | Bluepippin | 15 | 132,438 | 1,120,679,413 | 93.39 | WmBx15 | 123 | 386,404 | 94,855 | 74,581 | 134,608 | 11,667,272 | 2 |
| Universe | ELF | 15 | 119,121 | 599,671,617 | 49.97 | universeAx15 | 147 | 421,584 | 80,022 | 58,145 | 116,594 | 11,763,238 | 1 |
| Watchmaker Equinox | ELF | 15 | 89,058 | 419,979,074 | 35.00 | WmAx15 | 189 | 312,222 | 62,704 | 47,673 | 94,017 | 11,851,164 | 3 |
| SuperFi II | Bluepippin | 15 | 85,578 | 903,200,782 | 75.27 | superfiBx15 | 196 | 292,580 | 56,206 | 38,749 | 80,346 | 11,016,479 | 2 |
| Promega Go Taq Long | Bluepippin | 12 | 52,834 | 591,119,208 | 49.26 | promegaBx12 | 207 | 262,329 | 51,862 | 35,441 | 72,104 | 10,735,587 | 4 |
| SuperFi II | ELF | 15 | 96,073 | 536,211,718 | 44.68 | superfiAx15 | 217 | 262,033 | 51,414 | 33,211 | 76,546 | 11,157,036 | 2 |

For those amplification product libraries that gave >30x genome coverage low coverage index was calculated. The lowest LCI (indicating more even genome coverage was obtained with RepliQa followed by Terra polymerase (**Supplementary Figure 10**).

Long range PCR can often preferentially amplify smaller templates such that after multiple cycles the amplification reaction can be dominated by such shorter amplicons. Bluepippin size selected templates amplified by RepliQa and terra polymerase gave the longest average subread lengths (library insert size) of approximately 12kb (**Supplementary Figures 11 and 12**). With the larger 21kb ELF fractionated template the majority of reads were obtained from shorter amplification products. RepliQa gave the largest fraction of 20kb subreads (**Supplementary Figure 13**).

By comparing the PacBio HiFi data with the sequence of the *S. cerevisiae* S288C genome reference the error profile of the library generated after amplification with each enzyme could be determined. Terra, LongAmp and Promega Go Long are Taq based polymerase formulations and as a result were observed to give higher rates of particularly mismatch errors compared to the other enzymes that possess proofreading activity. NEB Q5 gave the lowest error rates (**Supplementary Figure 14**).

As might be expected those enzymes that gave the most even genome coverage also gave the best assembly statistics when 30x normalised coverage reads were assembled in the SMRTlink portal (**Table 2**). Here RepliQa followed by Terra polymerase gave the most contiguous assemblies. The *S. cerevisiae* genome is known to have 16 chromosomes [15] and additional circular chromosomal elements have been reported [16], therefore assemblies from material amplified using these enzymes has given near complete contiguity with the sum of contig lengths matching that expected for the yeast genome and with ELF fractionated fragments amplified for 15 cycles with RepliQa assembling into just 20 contigs.

## Discussion

It has been estimated that when sequencing the human genome 30x average genome coverage is required to give local base coverage at >15x so that both homozygous and heterozygous variants can be accurately detected [17] [18]. Indeed for Genomics Englands’ clinical human sequencing 95% coverage at >15 x is a key sequence dataset QC metric [19].

Here we compare sequence datasets from different PCR enzymes based on low coverage index, the percentage of the genome that is covered to <15x when using 30x datasets.

We have used 14 cycles PCR with 1ng of template in order to exacerbate any biases so to be able to differentiate between the enzymes used. Even with 14 cycles and a standardised input there was a broad range of yields. With those enzymes that gave a higher yield, fewer PCR cycles could have been used and indeed here we could have overamplified, possibly introducing bias. However, we obtained similar low coverage index values after 10 cycles PCR for these enzymes (results not shown).

The results presented demonstrate that there are distinct differences between PCR enzymes in the yield and evenness of amplification of fragments prior to sequencing. This further confirms that PCR can be a source of bias in genomic data and illustrates that the user should consider which enzyme is used for these applications, particularly for GC biased templates where coverage bias is more pronounced. Some enzymes work well, giving even coverage and low bias, in some situations, but few are capable of unbiased amplification of both GC and AT rich templates. Here, we have demonstrated that RepliQa, Watchmaker Equinox and Takara Ex Premier can amplify extremes of GC content with coverage bias similar to PCR free data and better uniformity than Kapa HiFi that we previously reported to give the best performance [4]. Surprisingly there was considerable variation in yield between enzymes, again illustrating that the user should carefully choose the enzyme used as utilising a low efficiency enzyme may give low yield, requiring the user to employ more cycles of PCR compared to other enzymes and accentuate the bias even more.

The performance of RepliQa, Equinox and also Takara Ex Premier reported here is impressive, with coverage uniformity after 14 cycles of PCR using 1ng template close to that obtained from PCRfree libraries made with 500ng input, even over wide extremes of GC content.

Here the genome of the malaria parasite has been a particular challenge. It is extremely AT rich with long stretches close to 100% AT [10]. Plasmodium causes a severe disease burden especially in sub Saharan Africa, in the 2021 World Malaria report there were 241 million malaria cases and 627 000 malaria deaths worldwide in 2020. As a result its genome is a major focus for researchers who often face challenges with human host DNA contamination and as those infected are mostly young children from which only low volume blood samples can be taken, only low amounts of DNA are often available. Using previously published modifications to cycling conditions we report extremely even genome coverage from just 1ng template using RepliQa, Watchmaker Equinox and Takara Ex Premier.

Sequencing of human genomes, particularly for variant discovery for academic and clinical outcomes, is a major application of NGS. The human genome is much larger and more complex than the microbial genomes used above and so the lead enzymes from this study were also tested for their ability to amplify limiting amounts of human genome template. DNA from the genome in a bottle standard NA12878 was used for this as it is probably the most sequenced human genome and has a good reference and validated list of variants [20]. Here the RepliQa, Watchmaker Equinox and Takara Ex Premier all performed remarkably well giving even coverage (low LCI) as well as high degrees of SNP and Indel precision all of which are better than the performance seen with Kapa HiFi. There are slight differences in performance with RepliQa and Watchmaker having greater SNP precision (**Table 1**) and RepliQa and Takara Ex Premier slightly better performance on Indels, meaning that from data generated here RepliQa would be the enzyme of choice as it gives superior performance both for Indels and SNPs.

In this study we have shown these enzymes to be capable of amplification of the complex genomes and genomes with extreme GC composition, to give uniformity of coverage similar to that obtained in PCR free library datasets. As a result we would expect these enzymes to perform well on other genomes irrespective of complexity and GC content, especially when using pure genomic DNA as used here. It should be noted that RepliQa is also reported to be tolerant to many known PCR inhibitors that can be present in crude extracts and has been shown to efficiently amplify targets from soil [21] and insects [22].

PCR amplification is also used ahead of long read sequencing either as a means of targeted sequencing or for low input template preparation. Standard ONT and PacBio library prep methods are PCR free and require over 1ug input DNA though both sequencing companies have protocols for low input library prep that involves amplification. We have compared a variety of polymerases for their ability to amplify 1ng of adapter ligated long template DNA and found the majority of enzymes to be quite inefficient for amplification of yeast genome fragments of 13.3 and 21kb and highlight a small number of enzymes that are capable of such reactions. The enzymes which performed best resulting in near complete genome coverage were RepliQa and Terra polymerase, though being a derivative of Taq Terra had higher rates of substitution error. Both these enzymes gave superior performance to Takara Primestar GXL reported the by Jia et al., 2014 to be the best enzyme for long template amplification [23], indicating that better enzymes are continuing to be developed.

Having better performing PCR enzymes will enhance the performance of NGS approaches, particularly for long read sequencing and yield the most contiguous assemblies and most complete analysis of genomic variants. At the time of this report RepliQa appears to give the best outcome for a variety of genomes and applications.

## Methods

### Genomic DNA

Human NA12878 CEPH/UTAH PEDIGREE 1463 DNA was purchased from Coriell Camden, NJ.

*Bordetella pertussis* Tohama I (ATCC-BAA-589D-5), *Escherichia coli* MG1655 (ATCC-700926D-5), *Clostridioides difficil*e 630(ATCC-BAA-1382DQ), *Plasmodium falciparum 3D7* (ATCC-PRA-405D) genomic DNA were obtained from ATCC via LGC standards, Teddington, UK.

*Saccharomyces cerevisiae* S288C genomic DNA (69240-3) was purchased from Merck, Gillingham, UK.

### Illumina Library construction

DNA (0.5µg in 100µl of 10mM Tris-HCl, pH8.5) was sheared in an AFA microtube using a Covaris S2 device (Covaris Inc.), with the following settings: for 200 bp fragments (duty cycle 20, intensity 5, 200 cycles/burst, 90 sec) or for 500 bp fragments (duty cycle 20, intensity 5, 200 cycles/burst, 30sec).

Sheared DNA was purified by binding to an equal volume of AMPure XP beads (A63881, Beckman Coulter, Inc. Brea, CA) and eluted in 50µl of 10mM Tris-HCl, pH8.5. End-repair, A-tailing and short Truseq adapter ligation* were performed using NEBNext UltraII (E7645L, New England Biolabs, Ipswich, MA). After ligation, excess adapters and adapter dimers were removed by Ampure XP clean-up with a 0.9:1 ratio of beads to sample, eluting in 50µl of 10mM Tris-HCl, pH8.5. Yield of adapter ligated fragments was ascertained by fluorimetric quantification using Qubit high-sensitivity DNA reagents (Q32851, ThermoFisher, Waltham, MA). *Oligo sequences are supplied in supplementary table S1.

PCR free libraries were prepared using the same approach starting with 500ng of genomic DNA input and ligating “IDT for Illumina” full length unique dual indexed adapters.

### PCR

Each PCR was performed using 1ng of template (Illumina truseq adapter ligated sheared DNA), 0.1nmol each of unique dual indexed i7 and i5 barcoding Illumina PCR primers (Supplementary table S1) and 25ul of 2x enzyme premix.

Unless indicated, all PCR reaction were performed for 14 cycles on an MJ Tetrad 4 thermocycler that had been validated for accuracy using a Driftcon thermocycler calibration instrument (Cyclertest, Landgraaf, Netherlands).

Cycling conditions used for each enzyme are detailed in **Supplementary Table 2,** with annealing at 60lJ for 15 seconds, and with denaturation and extension parameters as recommended by each manufacturer.

PCR reaction products were cleaned and size selected using a 0.7:1 ratio of AMPure XP SPRI beads (Beckman Coulter Inc.) to sample, according to the manufacturers’ protocol with elution in 30ul of EB buffer.

### Illumina sequencing

Prior to sequencing libraries were quantified by real-time PCR, using the SYBR Fast Illumina Library Quantification Kit (Kapa Biosystems cat. no. KK4834).

Libraries were pooled in an equimolar fashion whilst correcting for genome size to facilitate equal sequence coverage from each.

Samples were sequenced on an Illumina Novaseq 6000 instrument with 150 paired end read length and v1.5 chemistry.

### Long fragment amplification and Pacific Biosciences Sequencing

Two aliquots of S. cerevisiae DNA S288C were sheared using a Megaruptor 3 (Diagenode, NJ); aliquot 1: 5ug diluted to 150 ul volume with EB buffer was sheared at speed 30 and then 31 to tighten peak, aliquot 2: 10ug diluted to 310 ul volume with EB buffer was sheared at speed 31. Both aliquots of sheared DNA were purified and concentrated using a 1:1 ratio of PacBio AMPure SPRI beads with elution in 30ul EB. The amount of sheared DNA was measured by fluorimetry using Qubit high sensitivity DNA quantification reagents. Aliquot 1 had 3.1ug, and was size selected on Sage Sciences ELF instrument aiming for a maximum fragment size capture around 20kb. Aliquot 2 had 4.8ug, and 2.4ug was size selected on Sage Sciences Blue pippin instrument with selection set to range 10kb-50kb. Size selected fractions were purified using a 1:1 ratio of PacBio AMPure SPRI beads with elution in 50ul EB. Prior to amplification Illumina adapters were added to the ends of each fragment to enable primer annealing. End-repair, A-tailing and short Truseq adapter ligation were performed using NEBNext UltraII as above. After ligation, excess adapters and adapter dimers were removed by Ampure XP clean-up with a 0.9:1 ratio of beads to sample, eluting in 50µl of 10mM Tris-HCl, pH8.5. Yield of adapter ligated fragments was ascertained by fluorimetric quantification using Qubit high-sensitivity DNA reagents; ELF fractionated DNA was 10.2ng/ul, Blue pippin fractionated DNA was 3.04 ng/ul. Each was diluted to 1ng/ul with EB and 1ul of these dilutions used as template for long range PCR.

After PCR amplified products were concentrated with a 1:1 ratio of PacBio AMPure SPRI beads with elution in 7ul EB, 1ul of which was used for QC and 5ul for PacBio barcoded overhang adapter amplicon library prep with a different barcoded adapter being used for each successful PCR reaction [24].

PacBio libraries were quantified using Qubit high-sensitivity DNA reagents and where possible ligated samples were equimolar pooled, weak samples were pooled in their entirety. Pooled barcoded amplified fragments were sequenced using a Pacbio Sequel IIe instrument using binding kit v2.2 and sequencing chemistry 2.0, library was loaded at 80pM.

### Data processing and analysis

After sequencing, reads were mapped to each genome reference sequence using Minimap2 [25]. SAMtools [26] was then used to generate pileup and coverage information from the mapping output.

The quality of the sequence data was assessed using FastQC v0.11.9.

For human genome sequence data fastq files were automatically aligned to reference GRCh38 and were mapped using bwa version 0.7.17-r1188 and the command bwa mem -t 12 -p -Y -K 100000000 <reference.fa> <read1.fastq.gz>lJ<read2.fastq.gz> and duplicates were marked using biobambam2 version 2.0.79.

The resulting CRAM files were converted to BAM, sorted and indexed with samtools v.1.15.1. The stats and graphs were produced using samtools v.1.15.1.

All bams were subsampled to approx. 33X using samtools v 1.15.1 and seed 42. Variants were called using GATK HaplotypeCallerlJv3.5 with special options ‘- stand_call_conf 2 -stand_emit_conf 2 -A BaseQualityRankSumTest -A ClippingRankSumTest -A Coverage -A FisherStrand -A LowMQ -A RMSMappingQuality -A ReadPosRankSumTest -A StrandOddsRatio -A HomopolymerRun -A TandemRepeatAnnotator’[27].

For each dataset the overlap of variants on chromosomes 1-22 with the GIAB (Genome in a Bottle) v4.2.1 benchmark dataset was calculated using bcftools isec version 1.15.1. The true positive (TP) value was calculated as the number of variant sites that were identified in both the sample and and the GIAB benchmark dataset. The false negative (FN) was calculated as the number of variant sites identified in the GIAB dataset but not identified in the sample. The sensitivity was calculated as TP/(TP+FN) or the number variant sites found in the sample as a percentage of all the variant sites in the giab benchmark dataset [28].

PacBio HiFi read assembly was performed on 30x downsized coverage using IPA [29] assembler within SMRTlink v10.2.

## Abbreviations

NGS: Next-Generation Sequencing
Tm: Melting Temperature
bp: Base Pairs
PCR: Polymerase Chain Reaction
LCI: Low coverage Index.

## Declarations

### Ethics approval and consent to participate

Not applicable. The only human DNA used in this study was from a commercial source.

### Consent for publication

Not applicable

### Availability of data and materials

All datasets have been deposited in the ENA read archive. Human genome datasets under accession number ERP141249 (https://www.ebi.ac.uk/ena/browser/view/PRJEB56321), and microbial genomes under accession number ERP141224 (https://www.ebi.ac.uk/ena/browser/view/PRJEB56300).

## Authors’ contributions

MQ performed the experiments and performed primary data analysis. MQ designed the experiments. CC and JU performed the PacBio library prep and sequencing work. MQ wrote the manuscript. YGU and JK carried out bioinformatics analysis.

## Acknowledgements

The authors thank the Wellcome Trust Sanger Institute core sequencing and informatics teams.

Steven Leonard and IDT for design of barcode sequences.

